# Simple nanofluidic devices for high-throughput, non-equilibrium studies at the single-molecule level

**DOI:** 10.1101/201079

**Authors:** Carel Fijen, Mattia Fontana, Serge G. Lemay, Klaus Mathwig, Johannes Hohlbein

## Abstract

Single-molecule detection schemes offer powerful means to overcome static and dynamic heterogeneity inherent to complex samples. Probing chemical and biological interactions and reactions with high throughput and time resolution, however, remains challenging and often requires surface-immobilized entities. Here, utilizing camera-based fluorescence microscopy, we present glass-made nanofluidic devices in which fluorescently labelled molecules flow through nanochannels that confine their diffusional movement. The first design features an array of parallel nanochannels for high-throughput analysis of molecular species under equilibrium conditions allowing us to record 200.000 individual localization events in just 10 minutes. Using these localizations for single particle tracking, we were able to obtain accurate flow profiles including flow speeds and diffusion coefficients inside the channels.

A second design featuring a T-shaped nanochannel enables precise mixing of two different species as well as the continuous observation of chemical reactions. We utilized the design to visualize enzymatically driven DNA synthesis in real time and at the single-molecule level. Based on our results, we are convinced that the versatility and performance of the nanofluidic devices will enable numerous applications in the life sciences.

## INTRODUCTION

In many areas of the life sciences, single-molecule techniques are playing an increasingly important role in identifying and analyzing static or dynamic interactions of (bio-)molecules with high spatiotemporal resolution.^1–3^ Among the large number of available frameworks,^4^ single-molecule fluorescence detection (SMFD) schemes, such as confocal or camera-based total-internal-reflection fluorescence (TIRF) microcopy, are frequently chosen due to their simplicity, robustness and ease of use.^5^ However, users often have to choose between high throughput or high time resolution,^6–8^ many experiments require surface immobilization techniques to extend the observation time and it is difficult to access non-equilibrium conditions or follow reactions.

Many creative solutions have been proposed to overcome these hurdles using fluidic platforms^9,10^. These include mixers for studying single-molecule kinetics,^11,12^ titration devices,^13^ devices to confine molecules in polymer nanochannels,^14^ electroosmotic molecular traps,^15^ or microfluidic droplets containing individual enzyme analytes.^16^ The detection throughput can be increased by controlling the flow through the detection area; and, in the case of mixers, reactions can be triggered in front of the detection area. These fluidic platforms, despite their clear benefits, are however not yet widely used due to complex fabrication procedures, limited reusability and configurability or restrictions in integration into microscopy platforms.

Here, we introduce a novel fluidic platform for SMFD consisting of nano-/microchannel devices fabricated entirely in glass. Our platform uniquely combines several advantages: (1) Prolonged observation times of single analytes are achieved by geometrically confining the flow through nanochannels to a volume smaller than the excitation/detection focus along the long optical axis of a microscope. (2) High-throughput detection is possible by monitoring an array of nanochannels as well as by controlling the passage time through the nanochannels via the applied flow. (3) The devices are well-defined, robust and reusable. (4) The nanochannels can be functionalized to minimize, for example, non-specific adsorption of analytes. (5) The fluidic chips are compatible with a commercially available holder allowing a straightforward integration into a standard microscope stage as well as simple and reliable interfacing to tubing. (6) The flow rate can be controlled by cost-efficient syringe pumps. (7) Special nanochannel geometries can be used to continually mix and observe reactions at the single-molecule level.

We first demonstrate high-throughput sensing and tracking of short DNA oligonucleotides in parallel nanochannels utilizing single-molecule Förster resonance energy transfer (smFRET) between a donor and an acceptor fluorophore attached to the DNA. As our method allows for continuous detection, several hundred thousand events can be combined to gather reliable single-molecule data as well as obtaining sub-micrometer resolved velocity profiles within the nanochannels. We then demonstrate the mixing geometry (T-junction) by monitoring the conformational equilibrium of a DNA hairpin before, during and after mixing with a buffer containing a high salt concentration that stabilizes the closed hairpin conformation. Finally, we validate the potential of the device for monitoring enzymatic reaction kinetics by observing DNA synthesis at the single-molecule level.

## MATERIAL AND METHODS

### Fluidic device fabrication

Nano-/microfluidic devices were fabricated with a 45 mm × 15 mm footprint to be compatible with a commercially available chipholder Fluidic Connect PRO (Micronit Microtechnologies B.V., The Netherlands). Micronit also fabricated the devices. In brief, they consist of two thermally bonded borosilicate glass layers with photolithographically defined and wet-etched nano-and microchannels (Fig. 1). Microchannels and ports to connect to tubing are structured in the upper 1 mm thick glass, while nanochannels are etched into the bottom layer. Here, coverslip type D263 borosilicate glass (Schott) with a thickness of 175 μm was used enabling high-resolution fluorescence imaging with inverted oil-immersion microscope objectives with shortest working distances whilst minimizing undesirable autofluorescence.

**Fig. 1.**
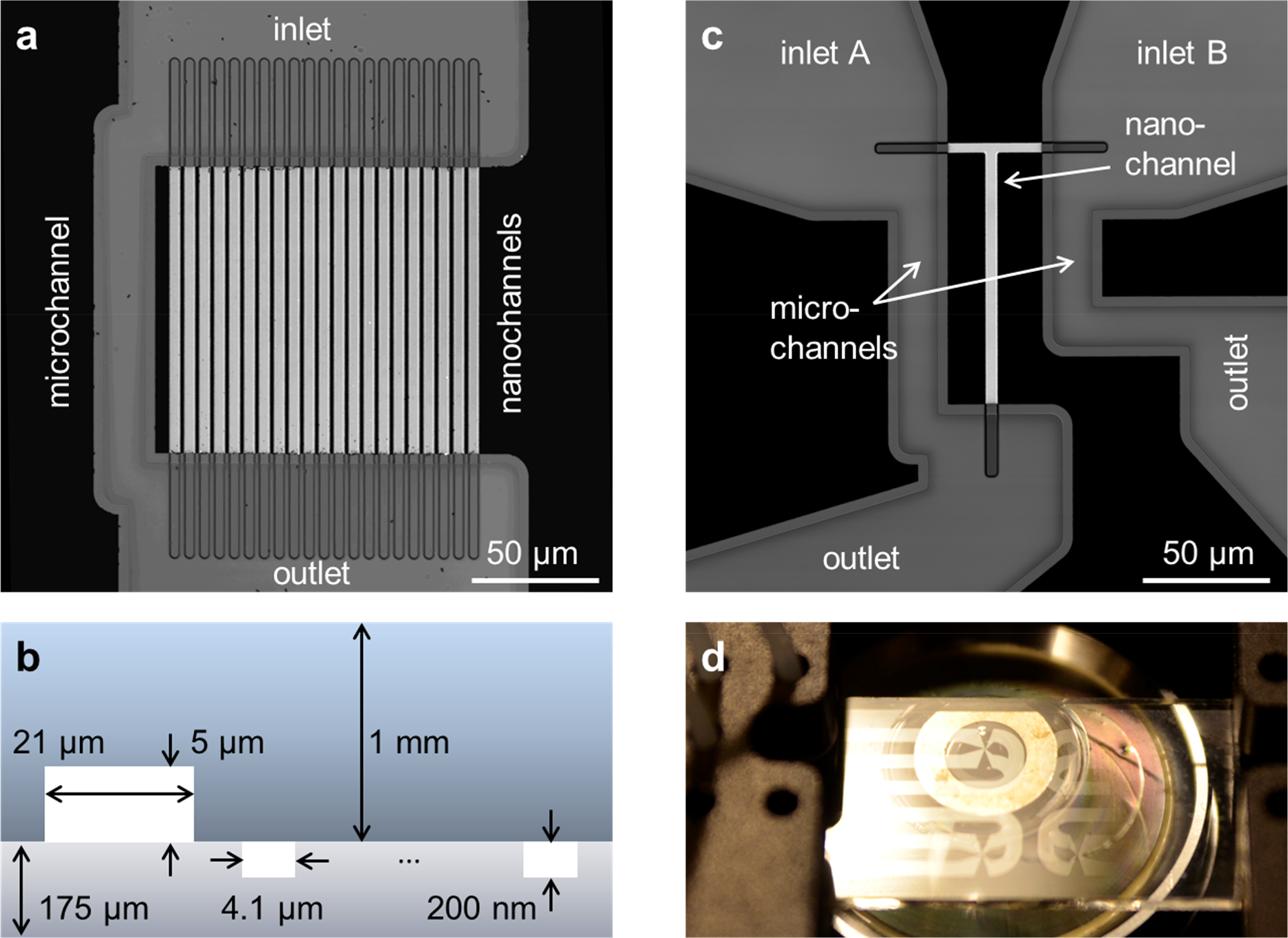
Design of nanofluidic chips. (a and c) High-resolution confocal scans based on reflection of light. (a) The parallel channel design contains 21 straight nanochannels (*l* × *w* × *h*: 120 by 4.1 by 0.2 μm) and a microchannel (*w* × *h*: 21 by 5 μm). (b) Schematic cross-section of the parallel channel array showing the dimensions (not to scale) of the microchannel etched into the top glass wafer and the nanochannels etched into the bottom glass wafer. (c) The design of the mixing device contains a single T-shaped nanochannel (horizontal part: *l* × *w* × *h*: 40 by 3.8 by 0.2 μm, vertical part *l* × *w* × *h* 100 by 4.7 by 0.2 μm) in between two microchannels. (d) Picture of a chip with 4 mixing channels, one of which is centered on our TIRF objective. Chips fit in a Micronit chip holder for easy connection to tubes and pumps (see also Fig. S2).

### Fluidic flow control

Flow was driven by Pump 11 Pico Plus Elite (Harvard Apparatus, USA) syringe pumps. For high nanochannel flow rates (50 pL/min) the pump generates a pressure of about 200 kPa. Due to the large ratio of flows in PFC, the dead volume in the 1 mm wide and up to 2 cm long feeding microchannels is replaced almost instantaneously. In PFC, low flow rates are achieved by dividing the syringe flow rate *Q* into the microchannel and array of nanochannels according to the channels’ hydraulic resistance, which is calculated as^17,18^

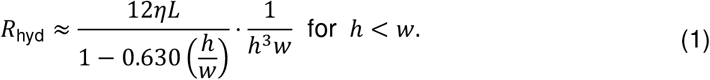

Here, *η* is the dynamic viscosity of 0.001 Pa s in water and *h*, *w* and *L* are the height, width and length of a nanochannel, respectively. (A pressure Δ*p* across a channel drops according to the Hagen Poiseuille Law *Q* = Δ*p*/*R*_*hyd*_, and for parallel resistances 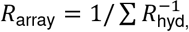 applies.) Due to the cubic dependence *h*^3^, the flow rate in each nanochannel is reduced by a factor of 60.000 compared to the microchannel with *L* = 140 μm, *w* = 10 μm, *h* = 5 μm compared to the applied syringe flow. Given the dimensions of the nanochannels, the flow is dominated by viscosity while inertia is negligible (Reynolds number <10^−4^, see Supporting Information Note 1); under these conditions laminar flow dictates the motion of the fluid and the equation for Poiseuille flow can be used to model the velocity field inside a nanochannel. In our design, the channel width is one order of magnitude larger than the height; as a consequence, the system resembles flow between two infinite parallel plates and the velocity is constant throughout most of the width (Supporting Information Note 2 and Fig. S5).

For the mixing device (Fig. 1c,d and Fig. S1b), two syringe pumps deliver flow to both feeding nanochannel inlets (each 20 μm long, 3.8 μm wide and 200 nm high). A single nanochannel (100 μm long, 4.7 μm wide and 200 nm high) is positioned downstream of the junction. Two bypassing microchannels lead to an overall reduction factor of 40.000 of the nanochannel flow compared to the combined syringe flow. Each feeding nanochannels has a hydraulic resistance of 25% of the long nanochannel, thus, a backflow into another feeding channel is prevented for differences in the syringe pump rate of up to 25%. When mixing fluidics of different viscosities (e.g., 1 M aqueous NaCl has a 10% higher viscosity than water^19^), pump rates can be conveniently adjusted to compensate for the different effective hydraulic resistances and to ensure a 1:1 mixing ratio.

### DNA

Fluorescently labelled oligonucleotides were ordered from IBA, Germany. To construct the DNA hairpin, a 30-mer primer sequence (biotin-5’-CCT CAT TCT TCG TCC CAT TAC CAT ACA TCC-3’) was annealed to a 75-mer hairpin sequence (5’-TGG ATT AAA AAA AAA AAA AAA AAA AAA AAA AAA AAA AAA TCC ATT GGA TGT ATG GTA ATG GGA CGA AGA ATG AGG-3’). The primer was internally labelled with ATTO647N at the -12 position; the hairpin was labelled with Cy3B at the 5’ end. Gapped DNA, used to study KF polymerase binding, was constructed using the same primer sequence, annealed to a template strand (5’-CCA CGA AGC AGG CTC TAC TCT CTA AGG ATG TAT GGT AAT GGG ACG AAG AAT GAG G-3’) and a downstream complementary strand (5’-TAG AGA GTA GAG CCT GCT TCG TGG-3’). The template strand was labelled with Cy3B at the +12 position. The DNA sensor used for the polymerization experiment consists of the same primer and template sequences, but with ATTO647N on the -7 position and Cy3B on the +25 position, respectively. dNTPs were ordered from Sigma-Aldrich / Merck, Germany.

### Buffers

DNA constructs (as well as DNA polymerases, if specified) were diluted in an imaging buffer containing 50 mM Tris HCl (pH7.5), 100 μg/mL BSA, 10 mM MgCl_2_, 5% glycerol, 1 mM DTT, 1 mM Trolox, 1% glucose oxidase/catalase and 1% glucose. Trolox is a triplet-state quencher and prevents fluorophore blinking. Glucose, glucose oxidase and catalase was used as an oxygen scavenger system to prevent premature photobleaching of fluorophores^20,21^. The concentration of gapped DNA was 1 nM and, if used, the concentration of KF was 10 nM. DNA hairpin concentrations were 500 pM in parallel channels and 1 nM in the mixing channel. (DNA hairpins were diluted in a similar imaging buffer without magnesium, but with additional NaCl as specified.) Prior to mixing, the concentration of DNA polymerization sensors was 1 nM; the concentration of KF was 5 nM, and the concentration of dNTPs was 200 μM each. For this polymerization experiment, we added neutravidin directly to the imaging buffer in a concentration of 0.6 μg/mL to block 5’ end of the biotinylated DNA primer. We found this prevents the formation of a low *E*\* state, the cause of which is probably binding of multiple KF polymerases to the same DNA molecule. The imaging buffer was applied to a 100 μL syringe (ILS, Germany), which was then connected to the nanofluidic device using ethylene tetrafluoroethylene (ETFE) tubing (1/16” outer diameter, 0.010” inner diameter) (IDEX, USA).

### Surface passivation

To prevent non-specific adsorption in experiments with proteins, channels were passivated with PEG using a variation of a method described previously^22^. Burning the glass to remove organic contaminations was only performed for cover slips. The fluidic devices were first flushed and incubated (3 × 5 min) with a 1:50 (vol/vol) Vectabond:acetone solution. All subsequent washing and passivation steps were performed by flushing the channels with ~100 μL of the respective solutions. PEGylated channels were filled with PBS and stored in a humid chamber at 4°C.

### Single molecule detection

We used a home built TIRF microscope and a fiber-coupled laser engine (Omicron, Germany) equipped with lasers of four different wavelengths (405 nm, 473 nm, 561 nm, and 638 nm). Laser intensities were independently controlled by a home-written LabVIEW program. Divergent light from the fiber output is collimated (f = 30 mm, Thorlabs, Germany) and focused by a second lens (f = 200 mm, Thorlabs, Germany) into the backfocal plane of a 100x NA 1.49 objective (Nikon, Japan). A polychroic filter and a multi-bandpass filter (Chroma, USA) prevented laser light from entering the emission path. A tube lens focuses the emission on an aperture, which reduces the effective field of view to a rectangle. Next, the light is spectrally split into a blue, a green and a red emission channel. The three beams are focused on an Ixon Ultra 897 emCCD (Andor, UK) with 512 by 512 pixels (maximum acquisition rate: 56 Hz at full frame and 100 Hz after cropping the frame to 343 by 256 pixels). In our configuration 1 pixel on the camera corresponds to a length and width of 112 nm in the sample plane. We use a Rapid Automated Modular Microscope (RAMM) system as a stage holder (ASI, USA), combined with motorized *x*,*y*-scanning stage and a *z*-piezo for focusing.

Molecules were excited with 130 mW (561 nm and 638 nm lasers) measured after the fiber output within the fluidic devices and with 30 mW (561 nm) and 15 mW (638 nm) for the surface immobilized experiments. A stroboscopic alternating-laser excitation (sALEX)^23^ scheme was used to reduce motion blur of diffusing molecules. Laser pulse widths were 1.5 ms (fluidic devices) and 3 ms (surface immobilized experiments) in a frame time of 10 ms. Green and red pulses were aligned back-to-back (Fig. 2a), so that particle movement between a green and a red frame is minimal. Shorter laser pulses, and the necessary higher laser powers, were found to cause rapid bleaching.

**Fig. 2.**
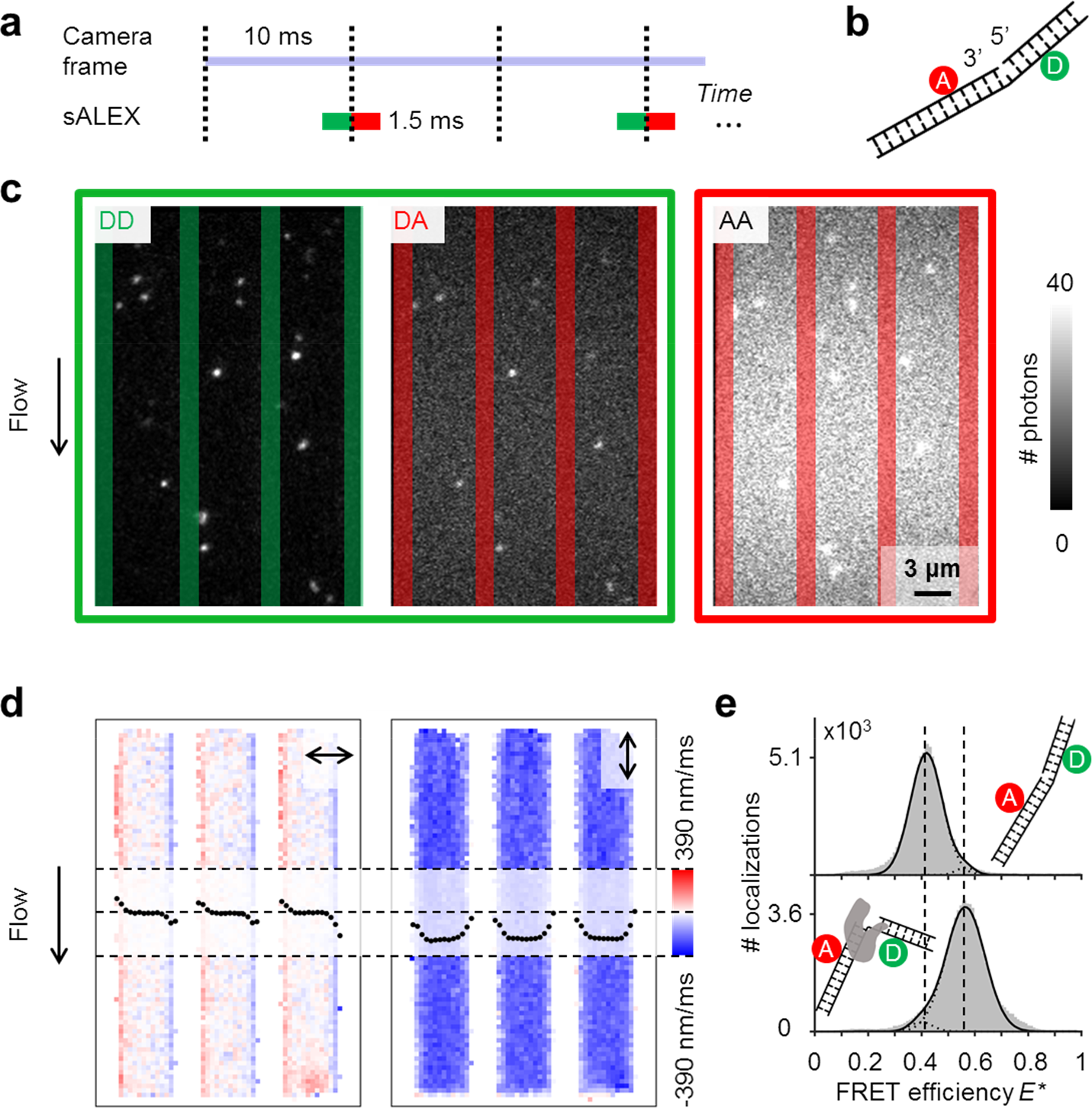
Single-molecule detection and flow profiles inside (parallel) nanochannels. (a) Schematic of stroboscopic alternating laser excitation (sALEX) in which the excitation time is considerably shorter than the required acquisition time of a camera frame. (b) Schematic of a gapped DNA construct labelled with a donor and acceptor dye located on opposite sides of the one-nucleotide gap. (c) Typical (non-averaged) movie frames showing our gapped DNA construct (*E*\*~0.4) flowing through the parallel nanochannels at a pump rate of 40 μL/h (Supporting Information Movie 2). Excitation colors are indicated by the surrounding boxes. For each molecule, photon counts of *DD* and *DA* after donor excitation are determined simultaneously. The photon counts of *AA* after acceptor excitation are collected during the next camera frame. Nanochannel boundaries are indicated with green bars (green detection channel) or red bars (red detection channel). Due to sALEX, motion blur is effectively suppressed and the back-to-back green and red excitation facilitates easy linking of *AA* to its corresponding signals *DD* and *DA*. (d) Flow velocimetry calculated from ~2 × 10^5^ *DA* and *AA* localization pairs perpendicular (left) and parallel (right) to the flow direction within the channels (binned 3 × 3 pixels, see also Material and Methods and Figs. S3 and S4). The resulting velocity profiles are superimposed on each map. (e) *E*\* histograms (100 bins) of our gapped DNA construct, measured in the parallel channels for 5 minutes. Top: 1 nM DNA. Bottom: 1 nM DNA in presence of 10 nM *E.coli* DNA Polymerase I (Klenow Fragment). Dashed lines are added for visual guidance. For full *E*\*/S histograms see Figure S7.

### Extracting emission intensities from movie frames

Particles were localized and tracked with a home-written variation on GaussStorm (Matlab).^24^ Time traces, histograms and binned maps were generated with custom-built software packages. The sALEX scheme effectively minimized motion blur and created mostly circular or elliptical intensity spots on the camera sensor. We applied a bandpass filter to find local intensity maxima before fitting the local maxima with elliptical 2D Gaussian functions from which we obtained the photon count as well as the position with sub-pixel accuracy.^24^ After filtering of localizations based on their intensities, our algorithm links corresponding particles in the green and red channels and calculates FRET efficiency *E*\* and stoichiometry ratio S for each molecule. A tracking algorithm is applied to track particles from frame to frame^25^.

### FRET, alternating laser excitation (ALEX) and accurate FRET

The apparent FRET efficiency *E*\* is calculated from the emission intensities of donor and acceptor after donor excitation (denoted *DD* and *DA*, respectively) for each molecule in each camera frame according to

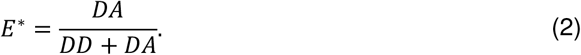

To verify the presence of an acceptor fluorophore on the DNA and to obtain the additional information required for correcting *E*\* for spectral cross-talk and detection efficiencies, we applied alternating laser excitation (ALEX) in which every frame of donor excitation is followed by a direct excitation of the acceptor fluorophore using a second laser resulting in a third photon stream (*AA*) for each molecule.^26, 27^ The detection of *AA* in addition to *DD* and *DA* allows for calculating the stoichiometry ratio S, defined as

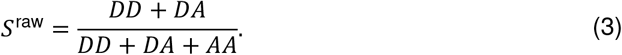

*S* can be used to filter molecules: molecules with a stoichiometry close to 0 have no photoactive donor (e.g., because of donor bleaching), and molecules with a stoichiometry close to 1 have no photoactive acceptor. Depending on the ratio of the laser intensities for direct donor and acceptor excitation, a stoichiometry around 0.5 represents molecules having both a photoactive donor and acceptor that can undergo FRET. In this work, we show FRET data based on molecules that have stoichiometry values 0.3 ≤ *S* ≤ 0.8 (nanochannel data) or 0.5 ≤ *S* ≤ 0.9 (immobilized hairpins).

ALEX also allows for easy correction of the apparent FRET efficiency *E*\* to obtain an accurate FRET efficiency E^27^. Corrections need to be made for 1) leakage of donor signal into the acceptor channel, 2) direct excitation of the acceptor when exciting the donor, and 3) quantum yields of the dyes and detection efficiencies of different colors on the camera sensor (summarized in the gamma factor). Leakage and direct excitation contributions can be deduced from the donor-only and acceptor-only peaks, respectively. Gamma will cause different FRET efficiencies to have slightly different stoichiometries; the contribution of gamma can therefore be deduced from the slope when plotting 1/*S* against *E*.

### Obtaining flow velocity profiles in parallel nanochannels

The displacements were calculated through localization and tracking of single molecules between consecutives frames. The travelled distance was calculated from sub-pixel localizations. The time between displacements was the time between the start of the green and the red excitation, respectively, i.e., 1.5 ms. Each calculated velocity was assigned to the starting position of the particle on the map. Then, the field of view was divided into regularly spaced bins of 3 × 3 pixels, so a bin will contain all the velocities of the particles that started in it. To avoid visualizing bins with only a few data points, a threshold was applied to exclude bins with <10 displacements from further analysis (Fig. S3). Excluded bins are located on the edges of the nanochannels.

## RESULTS and DISCUSSION

### Achieving parallel flow control

The device for high-throughput detection consists of an array of parallel nanochannels wet-etched into a first glass wafer (Fig. 1a,b, see also Material and Methods). Our array of nanochannels is connected in parallel via wafer-bonding with a bypassing larger microchannel etched into a second glass wafer such that the velocity of analytes in the nanochannel can be precisely tuned by the principle of parallel flow control (PFC) using a simple syringe pump.^28,29^ In PFC, nanochannel flow rates as low as 0.5–50 pL/min (corresponding to flow velocities of 10-1000 nm/ms) are achieved by dividing the flow into the microchannel and the array of nanochannels (see Fig. S1 for equivalent circuit diagrams) such that the flow rate in each nanochannel is reduced by a factor of 60.000 compared to the microchannel (Material and Methods).

Mixing devices have been primarily demonstrated for confocal applications, with latest designs even enabling two consecutive mixing steps^11,30^. Here, for the first time, we use the PFC principle for a mixing geometry: Two syringe pumps deliver flow to two microchannels and two respective parallel feeding nanochannel inlets which merge to a single nanochannel at a T-junction (Fig. 1c,d and Supporting Information Movie 1). PFC allows stable mixing conditions: the flow rates of both syringe pumps can differ by up to 25% before a parasitic backflow into one feeding channel occurs (Material and Methods).

### Detecting fluorescence emission of moving particles inside nanochannels

To characterize our devices, we monitored fluorescently labelled DNA molecules and their respective distance-dependent smFRET signatures. The apparent FRET efficiency *E*\* is calculated for each molecule in each camera frame using photon streams after donor excitation emitted by the donor and the acceptor (*DD* and *DA* respectively, see Material and Methods). To verify the presence of the acceptor, we applied alternating-laser excitation (ALEX) in which every frame of donor excitation is followed by a direct excitation of the acceptor fluorophores creating photon streams of *AA* (Material and Methods).^26,27^ Since motion blur severely affected our ability to determine emission intensities of individual molecules inside the nanochannels at frame times as short as 10 ms, we opted for stroboscopic ALEX (sALEX)^23^ in which the molecules are only excited for a fraction of the camera exposure time (Fig. 2a). We further aligned the green and red laser pulses back-to-back to facilitate the linking of a particle’s *AA* signal and position to its corresponding *DD* and *DA* signals from the previous frame.

### Obtaining velocity profiles in parallel nanochannels

We first measured 1 nM doubly labelled, gapped DNA (Fig. 2b) with a FRET efficiency of E*~0.4 for 10 minutes at pump rates of 40 μL/h. Upon donor excitation, both donor and acceptor intensities from individual DNA molecules are visible (Fig. 2c, left). Short excitation pulses (1.5 ms) lead to spots which are barely affected by motion blur and have an excellent signal-to-noise ratio allowing us to fit 2D Gaussians to obtain intensity numbers and sub-pixel localization accuracies for individual DNA molecules. The following direct excitation of the acceptor, provided information on the presence of the acceptor (Fig. 2c, right) and allowed us to calculate the displacements between the position of the molecule in the acceptor detection channel during donor excitation and during direct acceptor excitation.

Using 2 × 10^5^ displacements, we determined the particle velocities in vertical and horizontal direction and mapped the mean velocities back to the field of view; then, we constructed velocity profiles by averaging the mean velocities along the length of the channels (Fig. 2d). The velocity in the horizontal direction, perpendicular to the nanochannels’ axis, is expected to be zero as no flow is applied along this direction and diffusion is random; this is true in the center of the channels. For positions close to the walls, however, spatial restrictions to diffuse further left or right are translated into a net movement from the walls toward the center. The vertical velocity plateaus at −240 nm/ms in the center of each nanochannel are in excellent agreement with the theoretical model (Figs. S4 and S5). Along the nanochannels, the flow velocity profiles at different pump rates show a linear relationship between pump rate and flow speed (Fig. S4) consistent with laminar Poiseuille flow (Fig. S5).

We compared the magnitude of molecular movement due to advective flow with movement due to diffusion. To obtain typical values for the mean flow speed and diffusivity, we fitted all vertical displacements with a Cumulative Distribution Function (CDF)^31^ accounting for both flow and diffusivity, and applied it to a series of different pump rates (Fig. S6). We found an average diffusivity of *D* = 33 ± 1 μm^2^/s. The mean square displacement^32^ due to diffusion can be calculated using MSD = 2*D*Δt, in which Δt is the time between the beginning of green and red excitation, respectively (1.5 ms, due to back-to-back illumination). Using the diffusion coefficient *D*, we found a MSD of 0.1 μm^2^, giving a standard deviation of ~300 nm. CDF analysis also returned an average flow speed of 220 nm/ms, resulting in a flow-induced displacement of around 330 nm in 1.5 ms. These two displacements are in the same order of magnitude highlighting the fact that diffusion plays a major role at the time and length scales accessible in our experiments.

From our data, we compiled a FRET histogram with stoichiometry values 0.3 ≤ *S* ≤ 0.8 to select for doubly labelled DNA molecules (Fig. 2e). A single species is visible with a mean *E*\* of 0.4. To prove that our parallel device, after passivating the glass surface using PEG (see Material and Methods), is compatible with continuous enzyme detection, we added 10 nM of DNA polymerase I (Klenow fragment, KF) to the gapped DNA. We observed an increase in FRET efficiency from 0.4 to around 0.6, reflecting the shortened donor-acceptor distance upon KF binding and DNA bending (Fig. 2e).As mentioned above, we obtained ~2 × 10^5^ FRET data points within 10 minutes of measurement, with the current field of view on our setup (~29 μm by 19 μm) allowing to monitor 3 out of 21 channels. For comparison, in diffusion-based confocal smFRET experiments, fluorescence bursts of single molecules passing the focus are collected with a similar time resolution (~1-3 ms), but the time between individual bursts must be kept long enough to avoid doubly occupancy of the focal volume^7^. Assuming a time of 500 ms to obtain a single FRET data point, 10 minutes yield only around 1000 FRET data points; considerably less than in our nanochannels.

### Parallel nanochannels: resolving conformational dynamics of DNA hairpins at different salt concentrations

To probe the accessible dynamic range of our FRET measurements, we used DNA hairpins that can interconvert between an open (low FRET) and closed (high FRET) conformation. As reported before, the used DNA hairpin is mostly open at 0 M NaCl and mostly closed at 1 M NaCl, while showing fast opening and closing at an intermediate concentration of 0.5 M NaCl^23^. Our FRET histograms show that freely flowing DNA hairpins behave the same as immobilized ones, showing that we can access a broad FRET range (Fig. 3a,b). Here, the stroboscopic excitation scheme allows resolving the individual conformational states of “open” and “closed” despite the corresponding conformational dynamics being considerably faster than the frame rate of the camera. Individual time traces from freely flowing molecules show opening and closing of individual DNA hairpins in real time (Fig. 3c).

**Fig. 3.**
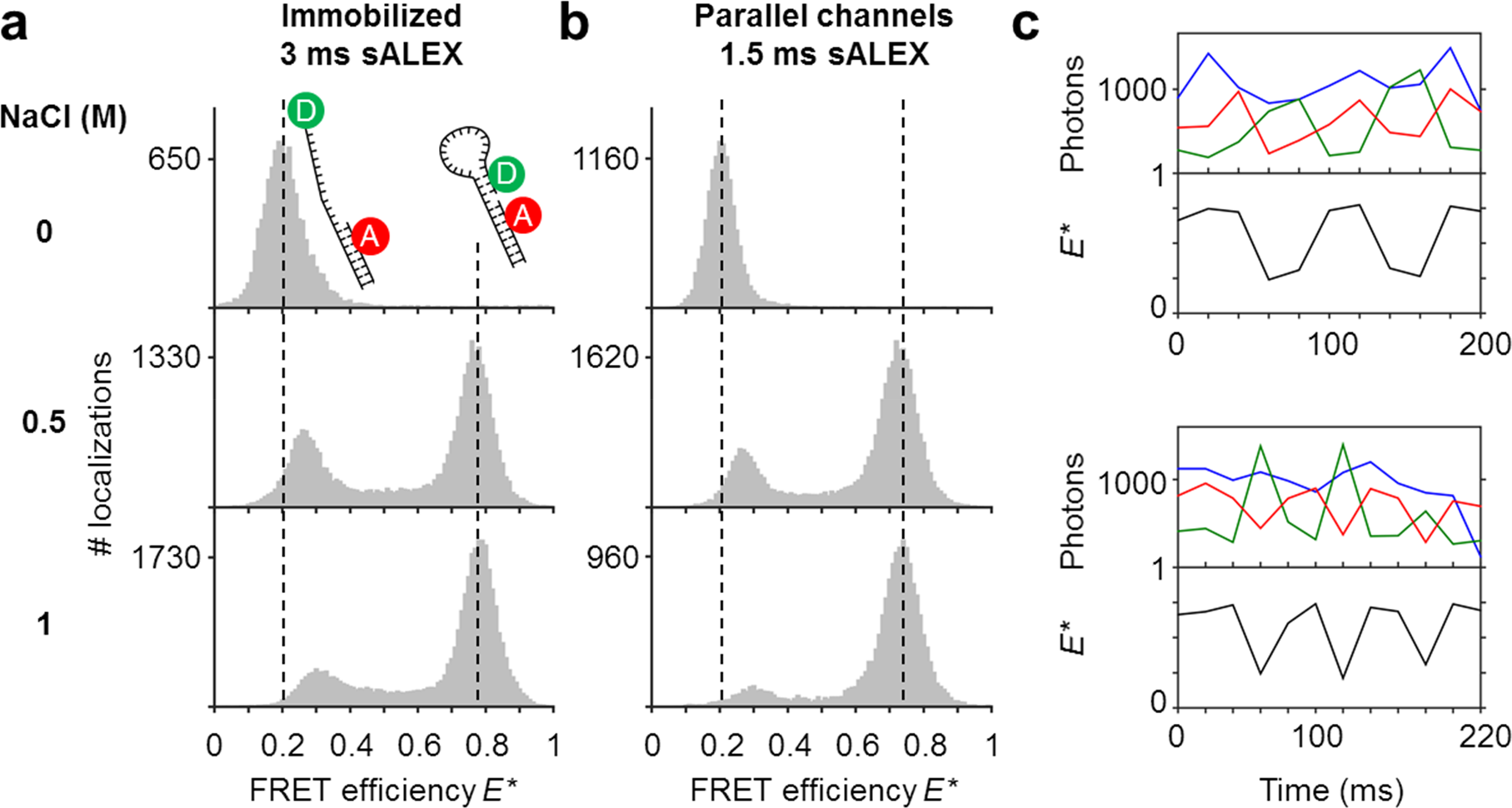
Single-molecule detection of DNA molecules immobilized (a) or freely flowing within parallel nanochannels (b). We analyzed the conformational equilibria of DNA hairpins as a function of the NaCl concentration and plotted FRET histograms (100 bins) derived from individual particle localizations. (a) DNA hairpins immobilized on a PEGylated coverslip are mostly open (low FRET) at [NaCl] = 0 M; the conformational equilibrium shifts towards the closed state (high FRET) with increasing salt concentrations. (b) The same DNA hairpin shows similar behavior when imaged in parallel nanochannels. When corrected for leakage, direct excitation and gamma, peak positions are the same as for the immobilized sample (Material and Methods and Supporting Information Table 1). For full *E*\*/S histograms see Figure S7. (c) Single-molecule time traces obtained from tracking individual molecules flowing through the channels at [NaCl] = 0.5 M. The anti-correlation of the *DD* (green trace) and the *DA* (red trace) signal and the resulting FRET time traces (black trace) indicate conformational changes in the millisecond timescale of the DNA hairpin during its passage. The *AA* signal (blue trace) remains constant.

### Nanofluidic mixing: triggering DNA hairpin closing

We used the mixing device to shift the conformational equilibrium from mostly open to mostly closed. To this aim, we mixed a buffer containing 1 M NaCl from the left with a buffer containing 0 M NaCl and DNA hairpins from the right. We constructed a map of binned hairpin localizations and corresponding FRET efficiencies (Fig. 4a). For each bin, we plotted the median FRET efficiency albeit the full spatially resolved FRET information remained preserved. For each of the three flow rate combinations depicted, a sudden shift from low to high *E*\* is observed around the junction. These positions mark the onset of mixing. As flow rates are increased, fewer hairpins diffuse far into the salt channel and the onset of mixing shifts slightly more towards the junction. Apparently, particle displacement due to diffusion, as opposed to particle velocity due to applied flow, plays a large role in mixing. In fact, in the nanochannel junction, mixing is purely diffusion-controlled and the mixing area becomes more confined as flow speeds increase.

**Fig. 4.**
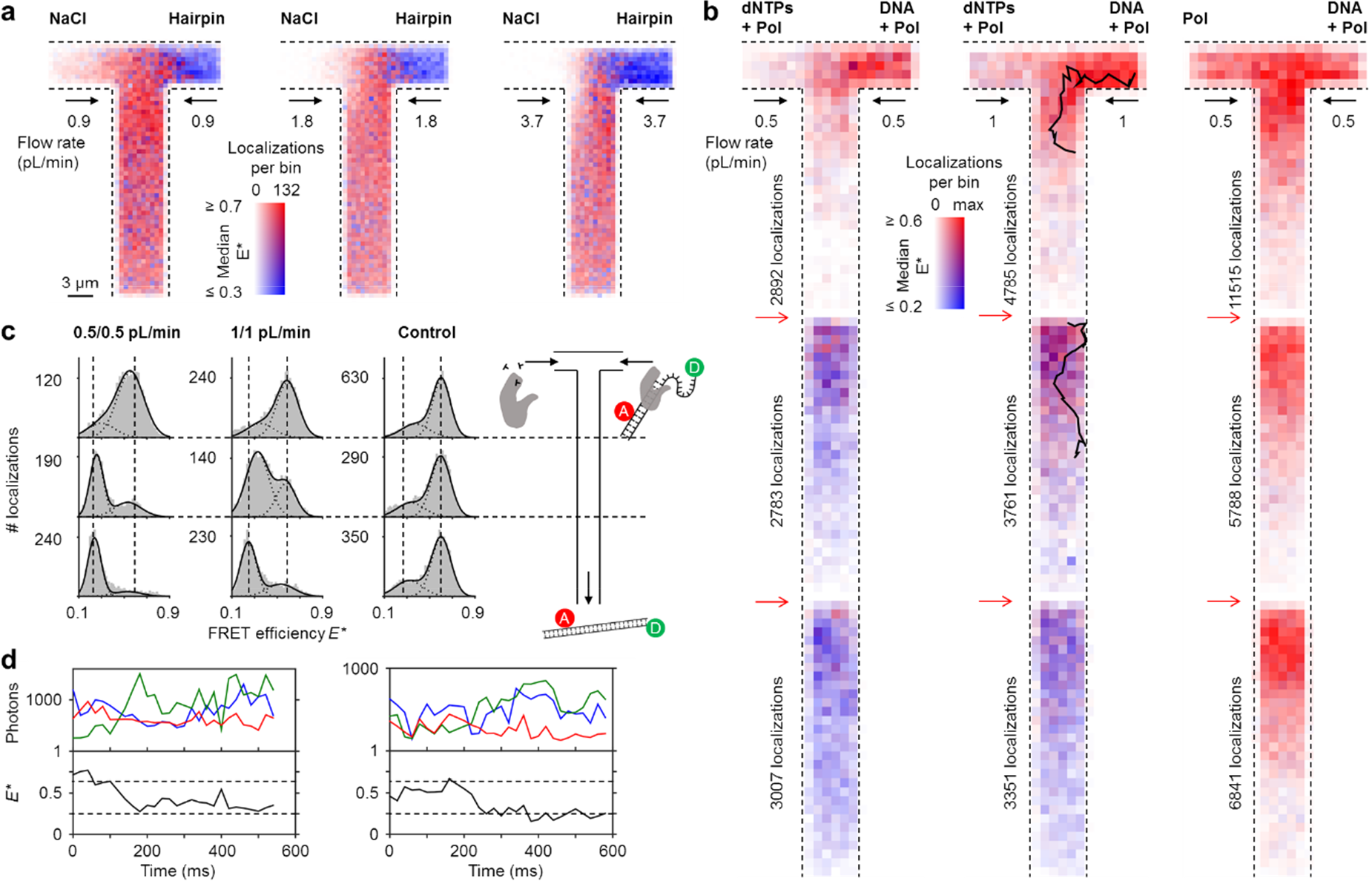
Mixing studies. (a) Mixing DNA hairpin (from the right, [NaCl] = 0 M) with a high salt solution ([NaCl] = 1 M) at different flow rates (Supporting Information Movie 3). The figure shows binned maps (4 × 4 camera pixels per bin), in which color represents the median FRET efficiency and opacity indicates the number of localizations in a bin over the time span of the measurement. (b-d) DNA polymerization (Supporting Information Movie 4). (b) Binned maps (8 × 8 camera pixels per bin) of mixing (DNA sensors + KF) from the right with (KF + dNTPs) from the left at two different pump rates. A control (outer right) did not contain dNTPs. Maps contain data from adjacent fields of view (sections); the borders of these sections are indicated with red arrows. Color represents the median FRET efficiency; opacity indicates the number of localizations in a bin, normalized for each map separately (maximum 58 for 0.5/0.5 pL/min, maximum 66 for 1/1 pL/min and maximum 126 for the control). Fewer molecules are localized in the lower parts of the channel sections because of fluorophore bleaching. (c) *E*\* histograms for each section of the mixing channel, as indicated with the schematic on the right. (d) Time traces of two molecules from the 1/1 pL/min condition. Upper panels: fluorescence intensities of *DD* (green), *DA* (red) and *AA* (blue). Lower panel: FRET efficiency *E*\* (black). Both traces show a decrease in FRET efficiency over time, suggesting polymerization of the DNA sensor.

This high diffusivity is advantageous for applications for which rapid mixing is required^33,34^ as turbulence does not occur in nanochannels due to very low Reynolds numbers. By combining the reduction of flow rates using PFC with the observation of single molecules, we established a precise control over diffusive mixing such that reagents can be detected before, during and after mixing, i.e. without any dead time.

### Nanofluidic mixing: triggering enzymatic catalysis of DNA synthesis

To establish our mixing device as a powerful tool for non-equilibrium studies, we utilized a FRET-based DNA polymerization sensor containing a primer DNA, labelled with an acceptor fluorophore, annealed to a long template DNA in which the single stranded overhang (25 bases) is labelled with a donor fluorophore (see Material and Methods).^35^ Since the single strand is coiled, the FRET efficiency is high (*E*\*~0.6) before decreasing substantially upon addition of DNA polymerases and nucleotides (dNTPs: deoxyribonucleotide triphosphates) to ~0.2. Using the same construct, we previously found that full polymerization takes on average 1.6 s when immobilized on a surface.^35^ In the nanofluidic device, we triggered the polymerization reaction by mixing (1 nM DNA + 5 nM KF) with (200 μM dNTPs + 5 nM KF) in a 1:1 mixing ratio, using low flow rates of either 0.5 pL/min or 1 pL/min (Fig. 4b). Similar to the DNA hairpin data at low flow rates (Fig. 4a), some molecules transiently diffuse into the other nanochannel. Due to the long reaction time, we acquired movies using three different fields of view and stacked them together to follow the reaction profiles until completion (Fig. 4b,c). The maps show a gradual decrease from high FRET efficiency (unpolymerized DNA sensor) near the junction to low FRET efficiency (polymerized DNA sensor) towards the outlet indicating that polymerization is completed within the time the molecules spend in the nanochannel: ~3 s at 1 pL/min and ~5 s for 0.5 pL/min nanochannel flow rates. This is on the same time scale as DNA polymerization of surface-immobilized DNA sensors. We note that at these low flow speeds, the fluorophores attached to the DNA molecules tend to bleach while being in the field of view due to the high laser intensities required to image molecules and the presence of proteins (KF). Two time traces indicating DNA synthesis in real time are shown (Fig. 4d).

In our current design, the distance from junction to outlet is 100 μm, corresponding to a residence time in the nanochannel of around 5 s using a flow rate of 0.5 pL/min and short DNA oligonucleotides. To gain access to further time points after mixing, designs using meandering channels could be implemented as demonstrated for confocal microscopy^11,30^ or widefield microscopy.^36^ Furthermore, our current field of view is cropped by a factor of two do ensure data acquisition at 100 Hz. With the use of faster cameras (sCMOS) and by reducing the overall magnification of the optical system (e.g., by replacing the 100x TIRF objective with a 60x objective), even more molecules could be simultaneously observed for a longer time over a larger area and with greater time resolution^37^.

## DISCUSSION

We pushed the boundaries of single-molecule detection under equilibrium and non-equilibrium conditions by introducing nanofluidic device designs that can be easily integrated in existing microscope setups: the first one with a series of parallel nanochannels for equilibrium studies, and the second with a single T-shaped nanochannel for non-equilibrium studies. For both cases, the core idea is the combination of camera-based single-molecule detection, which allows monitoring many fluorescent molecules in parallel, with constant replenishment of freely flowing molecules. Extended observation times are achieved by geometrical confinement of molecules, imposed by a nanochannel height of 200 nm. A stroboscopic excitation scheme resulted in near-circular emission spots on the camera sensor. Furthermore, in both designs we employed the concept of parallel flow control allowing us to work with simple syringe pumps whilst keeping the consumption of reagents low.

Our design using a parallel array of nanochannels is well suited for high-throughput measurements over extended periods of time. Moreover, longer tracks of single molecules flowing through the field of view can be observed by decreasing the sample concentration or decreasing the flow rate and increasing the viscosity of the buffer medium.

Using our nanochannel mixing devices, we accessed non-equilibrium conditions by mixing primarily open DNA hairpins from one inlet with a high-salt solution from a second inlet triggering hairpin closing, and showed that mixing is diffusive. Additionally, we observed polymerization of 25 bases on a DNA template by a DNA polymerase, illustrating that complex biological reactions can be followed in real time and in a continuous fashion.

We further note that the mixing device is conceptually equivalent to conventional stopped-flow instruments of the sort frequently used in the (bio-)chemical sciences to measure reaction kinetics of biomolecules, but with the unique benefit of allowing *continuous* observation of reactions. We therefore believe that the introduced nanofluidic designs represent a powerful platform for studying non-immobilized single molecules with high throughput and time resolution, under both equilibrium and non-equilibrium conditions.

## SUPPORTING INFORMATION

Supporting information is available online.

## ACKNOWLEDGEMENTS

We thank Timo Wenzel and Ebru Acun for initial characterizations of the fluidic devices. We thank Adrie Westphal for help with the confocal scans. J.H. acknowledges support from a Marie Curie Career Integration Grant (#630992). S.G.L. acknowledges financial support from The Netherlands Organization for Scientific Research (NWO) and the European Research Council (ERC) under Project 278801. M.F. acknowledges financial support from the Graduate School Experimental Plant Sciences (EPS), Wageningen University, under Project EPS3 3b 092. C.F. and M.F. contributed equally to this work.

## AUTHOR CONTRIBUTIONS

J.H. and K.M. conceived the project and designed the fluidic devices. C.F. performed the shown experiments. M.F. and J.H. performed preliminary experiments. C.F., M.F., and J.H. wrote and adapted software for data analysis and analysed the data. M.F. and K.M. performed fluidic simulations. All authors contributed in designing the research, participating in discussions and writing the manuscript.

